## Supplementary material for "Foam film vitrification for cryo-EM"

This document contains:

Extended discussion

Detailed procedure for foam film vitrification

Figs. S1 to S16

Table S1

Captions for Movie S1 to S15

References for SI reference citations

#### Guide for using foam film vitrification for cryo-EM sample preparation

##### Choosing surfactant

Among the surfactants tested in this study, some were unsuitable for the foam film method. LMNG, for example, showed a decreasing surface tension beyond its critical micelle concentration, but maintained a high tension (~50 mN/m) at 1% concentration (Fig. 3A). Further analysis revealed that LMNG's surface tension did not reach a minimum even at the maximum measurable bubble lifetime using our setup (20,000 ms) (Fig. 3B). This suggests LMNG forms micelles that are too stable to dissociate at the air-water interface, inhibiting foam formation. Similarly, GDN was unsuitable due to its high surface tension (~45 mN/m), while surfactants with very low water solubility, such as 12:0 PC and *E. coli* polar lipid extract had minimal impact on surface tension and failed to form stable foam films. Surfactants with a high critical micelle concentration can degrade image quality because the resulting abundance of micelles in the background lowers contrast (1). Some ionic surfactants, such as sarkosyl, can also be used for some stable specimens.

A practical approach to evaluate whether the surfactant is useful for foam film method is to dip the loop into a surfactant solution at the desired concentration. If a foam film forms and remains stable until its thickness drops below 100 nm, as indicated by the colour changes, the surfactant is suitable for this method. Stability can be further enhanced by adding a "control agent" such as glycerol. However, the glycerol concentration must be considered, as excessive amounts can reduce the signal-to-noise ratio in micrographs.

A method known as the shake test can also be used for surfactant screening, which involves measuring initial foam height and tracking its decay over time (2). Surfactants with good foamability and stability exhibit high initial foam heights and minimal decay. However, the shake test is limited by variability in the shaking power applied, as well as the simultaneous processes of foam generation and decay. More standardised methods, such as the Ross-Miles test or the Bikerman test, provide quantitative and reproducible results, allowing for a more precise evaluation of surfactant behaviour (3, 4).

##### Foam film thickness estimation

Foam film thickness can be visually estimated, without the need for laser interferometry, by observing interference colour patterns on the film. When the central region of the film within the loop becomes almost colourless leaving colour bands only at the lower half of the film, the thickness is approximately 100 nm in the centre. We found that this is the best moment for transfer. Because the thickness gradient across the film corresponds to an ice thickness gradient on the grid, we found that a foam film thickness of 60 – 90 nm prior to vitrification produces median ice thicknesses on grids ranging from 30 to 60 nm, good for imaging a wide range of macromolecules.

### Detailed procedure for foam film vitrification

1. **Materials**
   1) Solutions
      - Protein sample: e.g. at 5 g/L in 0.01% DDM (must adjust concentration and surfactant as needed).
   2) Materials and instruments
      - Liquid nitrogen (filtered)
      - Ethane (for cryocooling)
      - Loop: copper loop with an outer diameter of ~5.8 mm and thickness of ~0.15 mm (Fig. 1B) (adjust loop material and size as needed).
      - Container (PMMA): used for holding the sample solution and immersing the loop, plus a locking holder (Fig. 1E) (adjust container material as needed).
      - EM grids: your choice of grid type
      - Grid box: for storing vitrified grids
      - Tweezers (Dumont L5)
      - Manual plunger
      - A cryostat and temperature control unit (5)
      - Plastic beaker: to cover the cryostat and protect from air contaminants
      - Personal protective equipment: Face mask/shield, cryo gloves, safety glasses/goggles, lab coat
   3) Safety notes
      - Ethane is extremely flammable and cryogenic, and causes serious cold burns easily. Use in a well-ventilated area, wear appropriate cryo-protective gloves, and avoid sparks or open flames.
      - Liquid nitrogen can cause cold burns and when handled in large quantities asphyxiation in poorly ventilated areas; handle with care.
      - General: Always wear a lab coat, safety goggles/face shield, and suitable gloves when working with cryogens.

2. **Foam film vitrification procedure**
   Tip: Perform all steps at 4 °C if possible (e.g., in a cold room), to reduce evaporation and slow down foam film thinning.
   1) Cryogen setup
      - Pour filtered liquid nitrogen into the ethane container until full. Also fill the outer cup (surrounding reservoir) until full.
      - Cover the assembly with a plastic beaker or lid to protect from air contaminants.
      - Wait about 3 minutes for the liquid nitrogen to boil off (it will cool the container).
   2) Top up liquid nitrogen
      - Replenish liquid nitrogen in the outer cup so that it completely covers any grid boxes or holders.
      - By now, liquid nitrogen in the inner ethane container should have mostly evaporated, leaving the container cold but free of excess liquid nitrogen.
   3) Fill the ethane cup
      - Carefully condense ethane gas into the cold cup.
      - Follow your lab's safety protocols for handling flammable, cryogenic ethane.
   4) Set temperature
      - Transfer the cryostat to the manual plunger and connect to the temperature control unit.
      - Wait until the ethane temperature reaches about -180 °C (or your desired set point).
   5) Prepare grid box
      - Place dry, moisture-free grid boxes (with the cap screw loosened) into the grid box holder, ensuring they are chilled by the liquid nitrogen.
   6) Pick up an EM grid
      - Using tweezers, take an EM grid (carbon/gold side facing toward the sample application).
      - Mount the grid in the tweezer holder.
   7) Bend the grid
      - Gently bend the grid by ~5–15° so that the foam film can be applied nearly parallel to the grid surface (Fig. 1A).

- Caution: Excessive bending can cause grid squares to break.

8) Position the grid above the ethane surface
   - Hold the grid 50 to 100 mm above the ethane surface.
   - If it's too low, vitrification may be imperfect; if it's too high, the force upon contact ethane surface can break grid squares.
9) Prepare the sample container
   - Fill the PMMA container with the protein and surfactant solution.
   - Cover the container with its lid when not in use to minimise evaporation.
10) Immerse and pull the loop
    - Submerge the copper loop in the solution, then slowly pull it out.
    - Pulling speed affects initial film thickness: slower pulling results in thinner film.
11) Observe the colour bands
    - The thin film in the loop shows colour bands.
    - Position yourself so the light reflects off the film, making these bands visible.
12) Wait for film thinning (in the order of tens of seconds to few minutes)
    - Watch for the top half of the film to become transparent while the lower half still shows colour bands.
    - Thinning speed varies with humidity and solution viscosity.
13) Transfer the film to the grid
    - Once the top half is transparent, swiftly touch the middle region of the foam film.
    - The film should transfer onto the grid surface.
14) Plunge-freeze
    - Immediately plunge the grid into the liquid ethane at -180 °C.
    - The rapid freezing vitrifies the sample.
15) Store the grid
    - Using tweezers, transfer the vitrified grid into the grid box in liquid nitrogen.
    - Make sure the grid is fully submerged in liquid nitrogen to prevent devitrification.
16) Repeat for additional grids
    - Return to Steps 10–15 for each additional grid.
17) Finalize storage
    - Once done, securely close the grid box(es).
    - Transfer them to a liquid nitrogen storage dewar for long-term storage.

**3. Troubleshooting tips**

1) Film collapses too quickly
   - Increase humidity or decrease surfactant concentration.
   - Work at 4 °C to slow evaporation.
2) Grid squares break frequently
   - Reduce the bending angle.
   - Move the grid closer to the ethane surface or approach it more gently.
3) Ice too thick
   - Wait longer for the foam film to thin before transferring to the grid.
   - Wait after transfer the foam film to grid, however, the waiting time needs to be optimised.

#### References


1. C. Le Bon, B. Michon, J. L. Popot, M. Zoonens, Amphipathic environments for determining the structure of membrane proteins by single-particle electron cryo-microscopy. *Q Rev Biophys* **54**, e6 (2021).
2. O. Bartsch, Über Schaumsysteme. *Kolloidchemische Beihefte* **20**, 1-49 (1924).
3. J. Ross, G. D. Miles, An apparatus for comparison of foaming properties of soaps and detergents. *Oil & Soap* **18**, 99-102 (1941).
4. J. J. Bikerman, *Foams*, Applied physics and engineering v. 10 (Springer-Verlag, New York, 1973), pp. viii, 337 p. : illus.
5. C. J. Russo, S. Scotcher, M. Kyte, A precision cryostat design for manual and semi-automated cryo-plunge instruments. *Rev Sci Instrum* **87**, 114302 (2016).

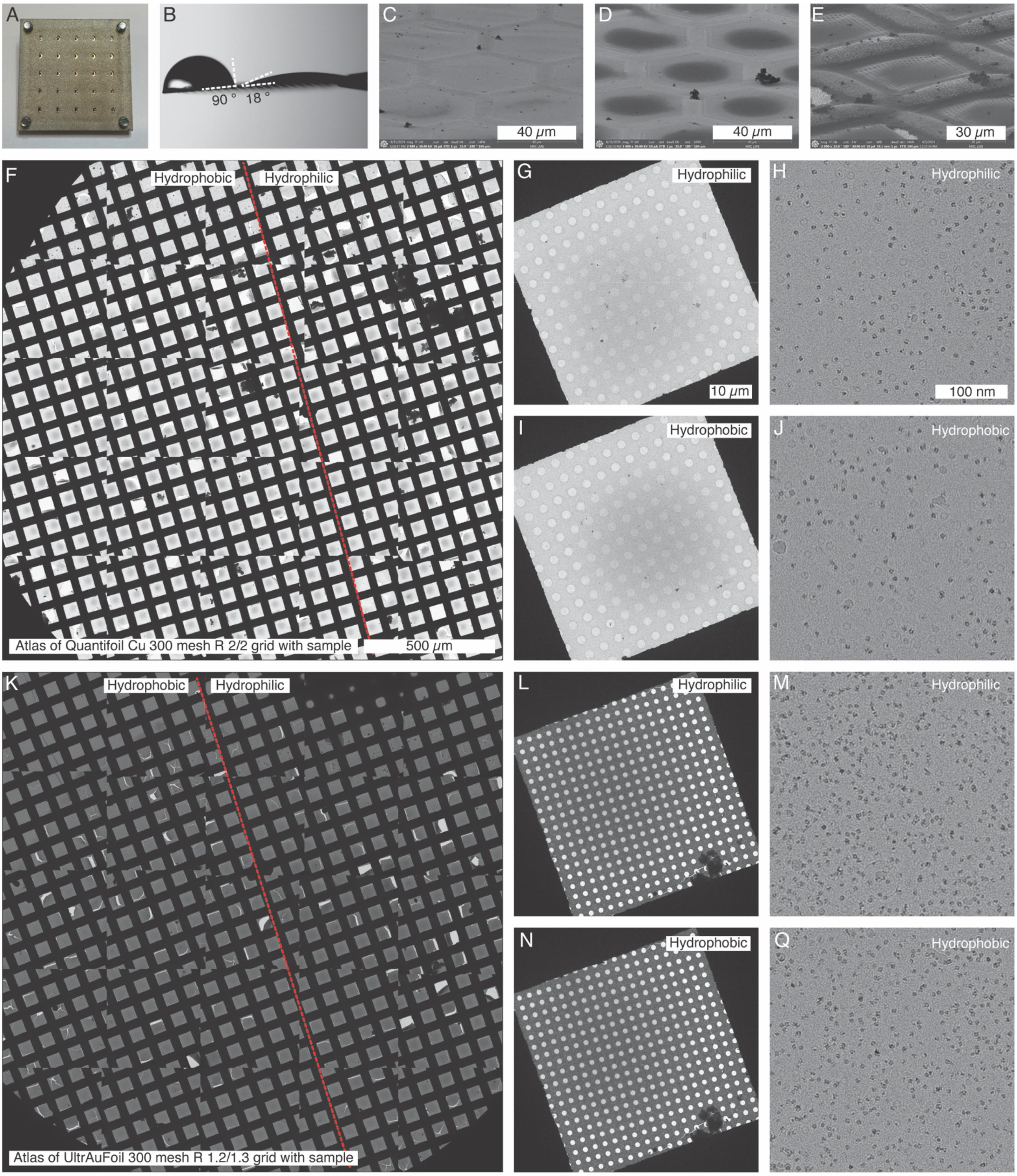


**Figure S1.** Evaluating the effect of grid wettability using the foam film method. (A) Grid holder with a mask covering half of the grid for glow-discharging. (B) Measurement of water contact angle on a half-glow-discharged grid. Scanning electron microscope images of (C) HexAuFoil grid with thin ice, (D) HexAuFoil grid with thicker ice, and (E) Quantifoil Cu 300 mesh R 2/2 grid with ice. (F) Atlas of a half-glow-discharged Quantifoil Cu 300 mesh R 2/2 grid, with the area to the right of the red dashed line glow discharged before foam film application. Images from the hydrophilic region include (G) a grid square and (H) a micrograph of a hole with horse spleen ferritin molecules visible. Images from the hydrophobic region include (I) a grid square and (J) a horse spleen ferritin micrograph. (K) Atlas of a half-glow-discharged UltrAuFoil 300 mesh R 1.2/1.3 grid, with the area to the right of the red dashed line glow discharged before foam film application. Images from the hydrophilic region: (L) a grid square and (M) a horse spleen ferritin micrograph; images from the hydrophobic region: (N) a grid square and (Q) a horse spleen ferritin micrograph.

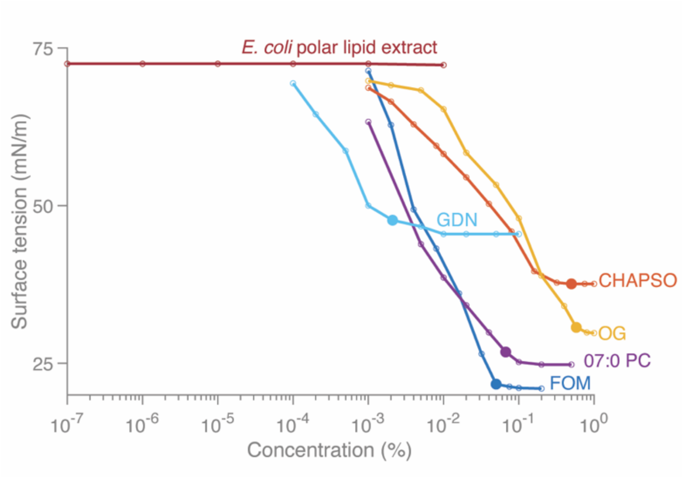


**Figure. S2.** Dynamic surface tension of FOM (blue), CHAPSO (orange), OG (yellow), 4) PC (purple), GDN (cyan) and *E. coli* polar lipid extract measured (red) at 18°C using the maximum bubble pressure method. Filled dots indicate the critical micelle concentration of each surfactant. The bubble generation time was set to 20 seconds to approximate equilibrium surface tension.

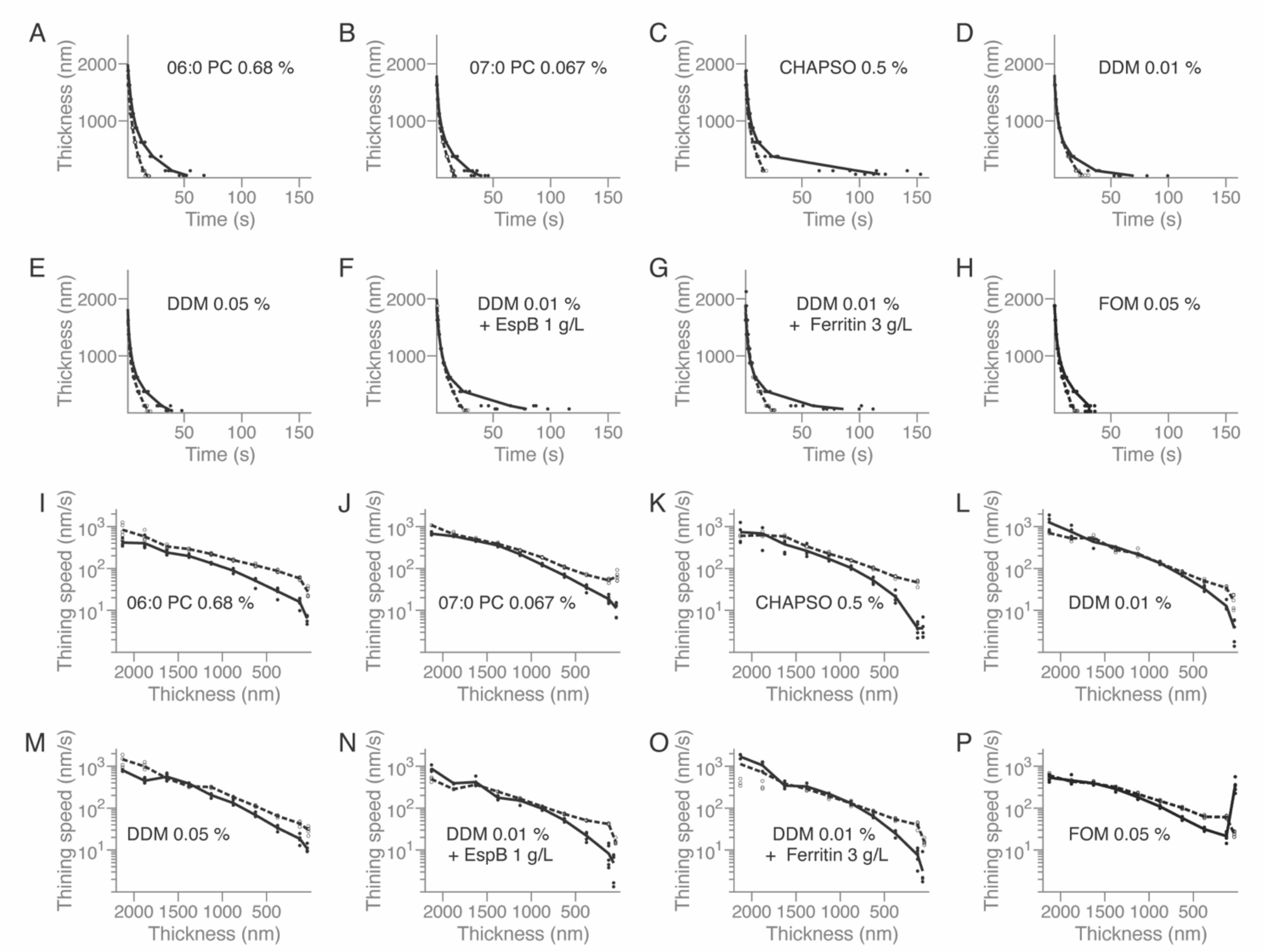


**Figure S3.** (A - H) Time-dependent changes in film thickness for various surfactant solutions mixed with different proteins at relative humidities of 60% (dashed curves) and 95% (solid curves) at 18°C. Dots represent individual measurements at specific thicknesses. (I - M) Speed of film thinning versus film thickness for different surfactant solutions mixed with various proteins at 60% (dashed curves) and 95% (solid curves) humidity levels at 18°C. Dots represent individual thinning speeds measurement at specific thicknesses.

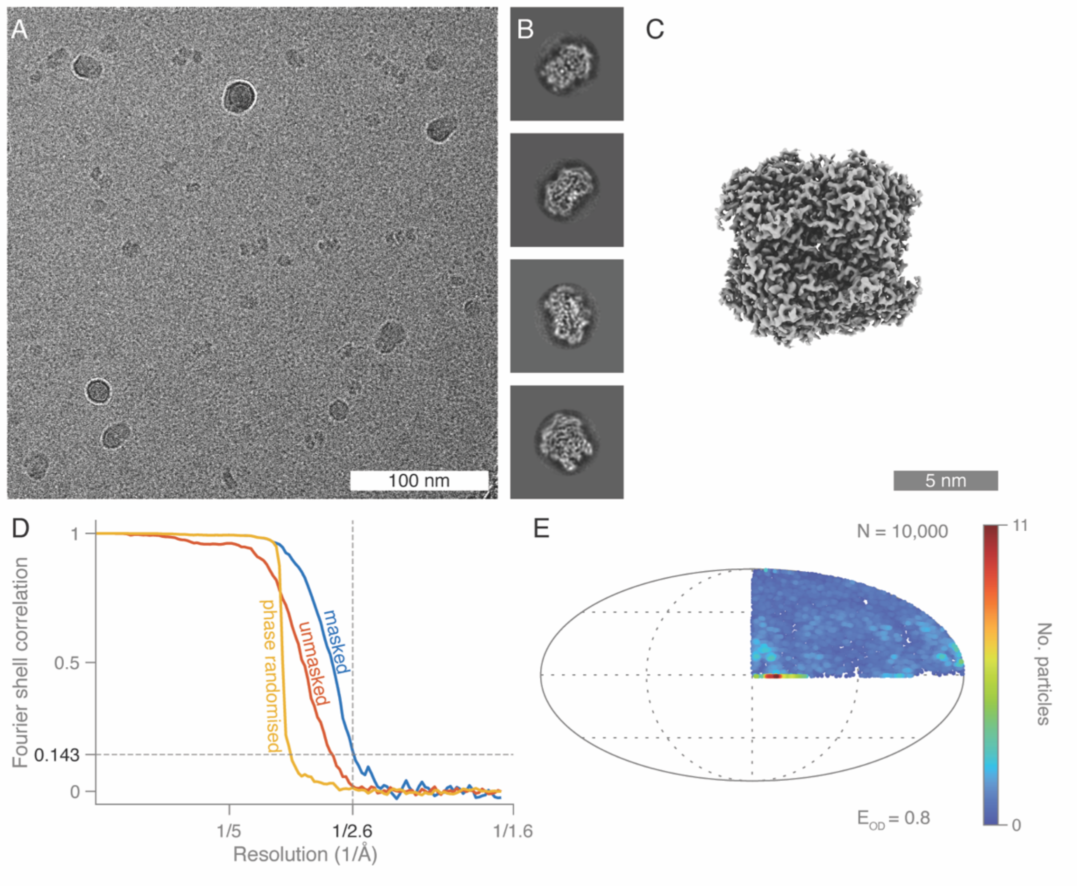


**Figure S4.** Structure determination of catalase with 0.01% DDM. (A) Representative cryo-EM micrograph. (B) Four selected 2D class averages. (C) Final reconstructed cryo-EM 3D map. (D) Fourier shell correlation curves: unmasked dose-weighted half-maps (red), phase-randomised dose-weighted half-maps (yellow), and final, independently refined, masked, dose-weighted half-maps (blue). (E) Asymmetric particle orientation distribution plotted on a sphere, with the orientation distribution efficiency indicated.

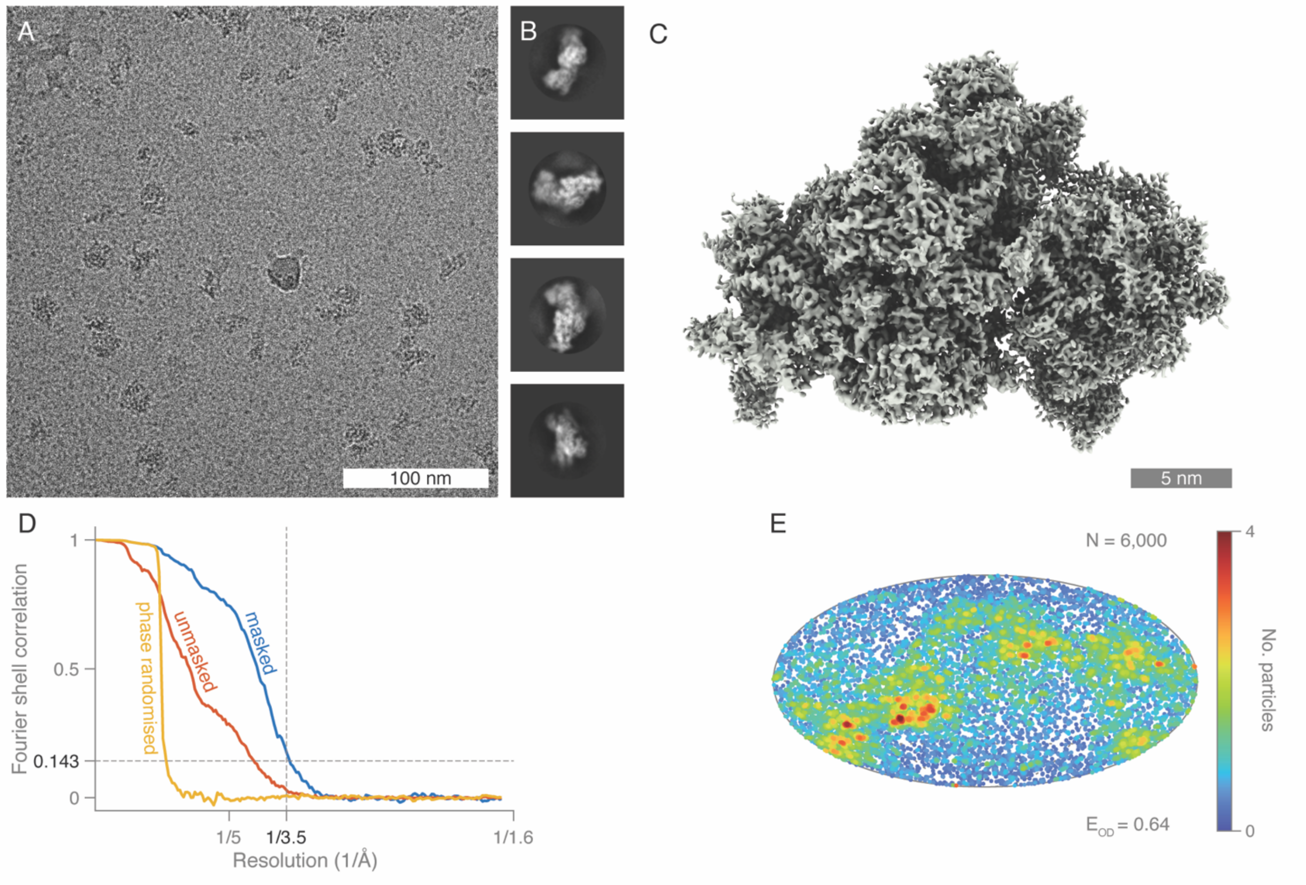


**Figure S5.** Structure determination of 30S ribosomes with 0.01% DDM. (A) Representative cryo-EM micrograph. (B) Four selected 2D class averages. (C) Final reconstructed cryo-EM 3D map. (D) Fourier shell correlation curves: unmasked dose-weighted half-maps (red), phase-randomised dose-weighted half-maps (yellow), and final, independently refined, masked, dose-weighted half-maps (blue). (E) Asymmetric particle orientation distribution plotted on a sphere, with the orientation distribution efficiency indicated.

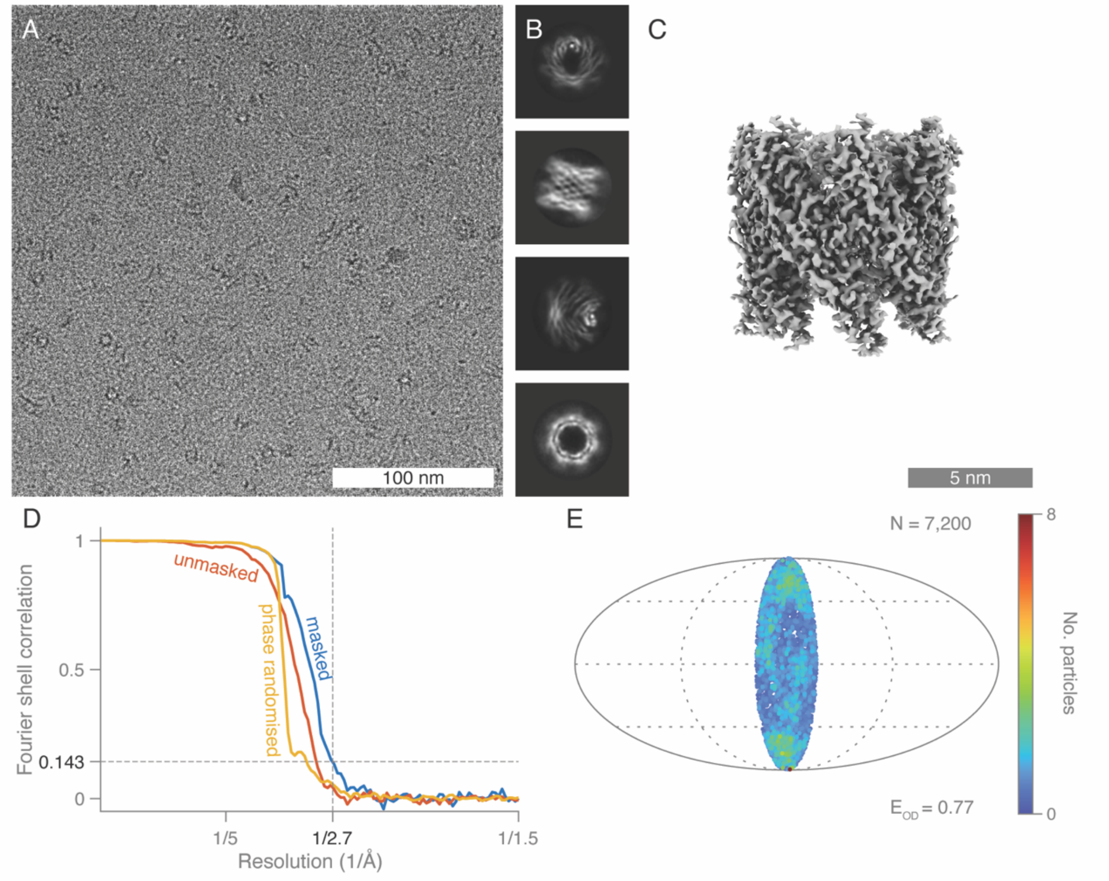


**Figure S6.** Structure determination of EspB with 0.01% DDM. (A) Representative cryo-EM micrograph. (B) Four selected 2D class averages. (C) Final reconstructed cryo-EM 3D map. (D) Fourier shell correlation curves: unmasked dose-weighted half-maps (red), phase-randomised dose-weighted half-maps (yellow), and final, independently refined, masked, dose-weighted half-maps (blue). (E) Asymmetric particle orientation distribution plotted on a sphere, with the orientation distribution efficiency indicated.

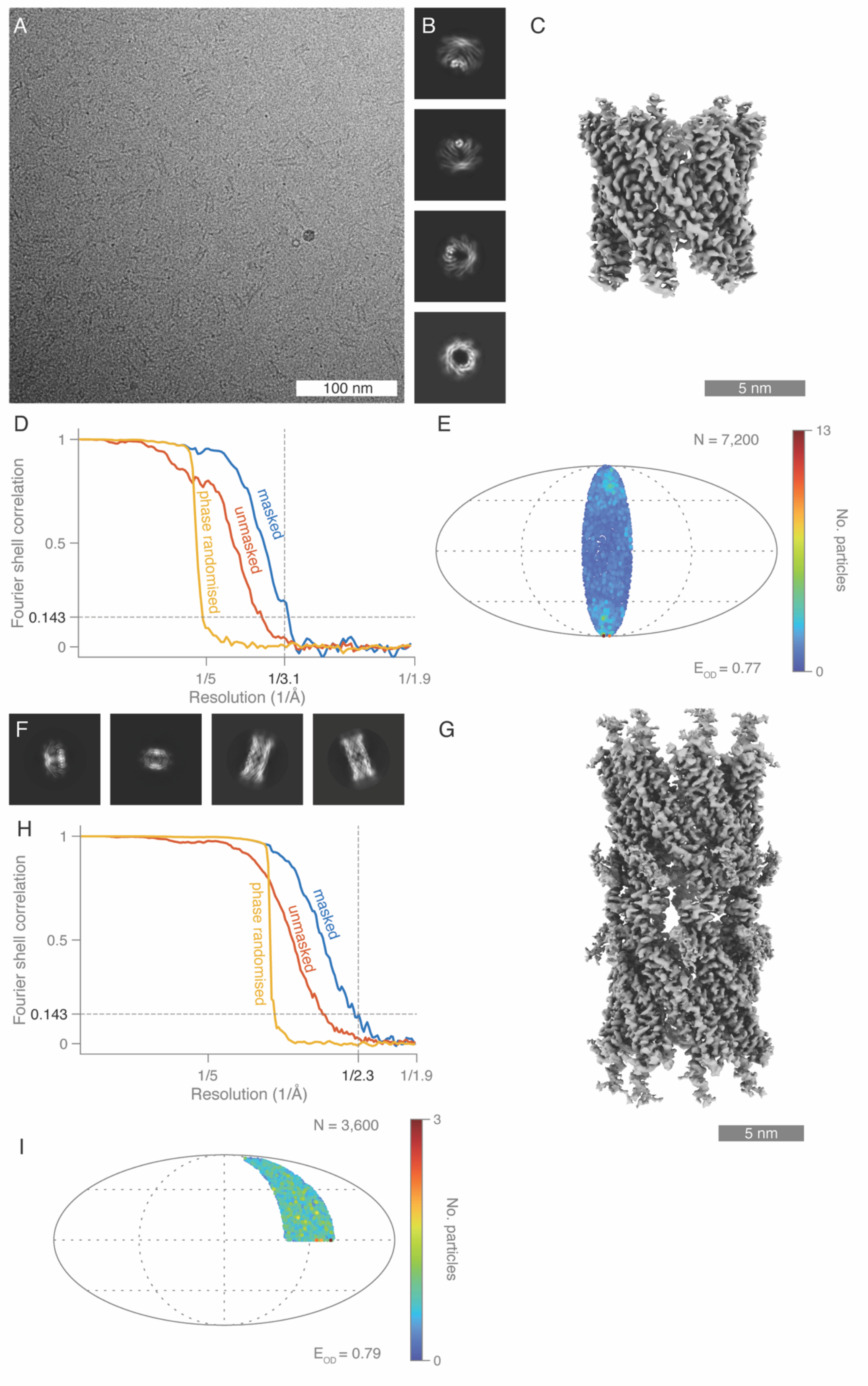

A
100 nm
B
C
5 nm
D
Fourier shell correlation
1
0.5
0.143
0
masked
unmasked
phase randomised
1/5
1/3.1
1/1.9
Resolution (1/Å)
E
N = 7,200
13
0
No. particles
E_OD = 0.77
F
G
5 nm
H
Fourier shell correlation
1
0.5
0.143
0
masked
unmasked
phase randomised
1/5
1/2.3
1/1.9
Resolution (1/Å)
I
N = 3,600
3
0
No. particles
E_OD = 0.79

**Figure S7.** Structure determination of EspB and EspB 14-mers with 0.67% 06:0 PC. (A) Representative cryo-EM micrograph of EspB. (B) Selected 2D class averages for EspB. (C) Final reconstructed cryo-EM 3D map of EspB. (D) Fourier shell correlation curves: unmasked dose-weighted half-maps (red), phase-randomised dose-weighted half-maps (yellow), and final, independently refined masked dose-weighted half-maps (blue). (E) Asymmetric particle orientation distribution represented on a spherical plot. (F) Selected 2D class averages for EspB 14-mers. (G) Final, reconstructed cryo-EM 3D map of EspB 14-mers. (H) Fourier shell correlation curves for EspB 14-mers: unmasked dose-weighted half-maps (red), phase-randomised dose-weighted half-maps (yellow), and final, independently refined masked dose-weighted half-maps (blue). (I) Asymmetric particle orientation distribution plotted on a sphere, with the orientation distribution efficiency indicated.

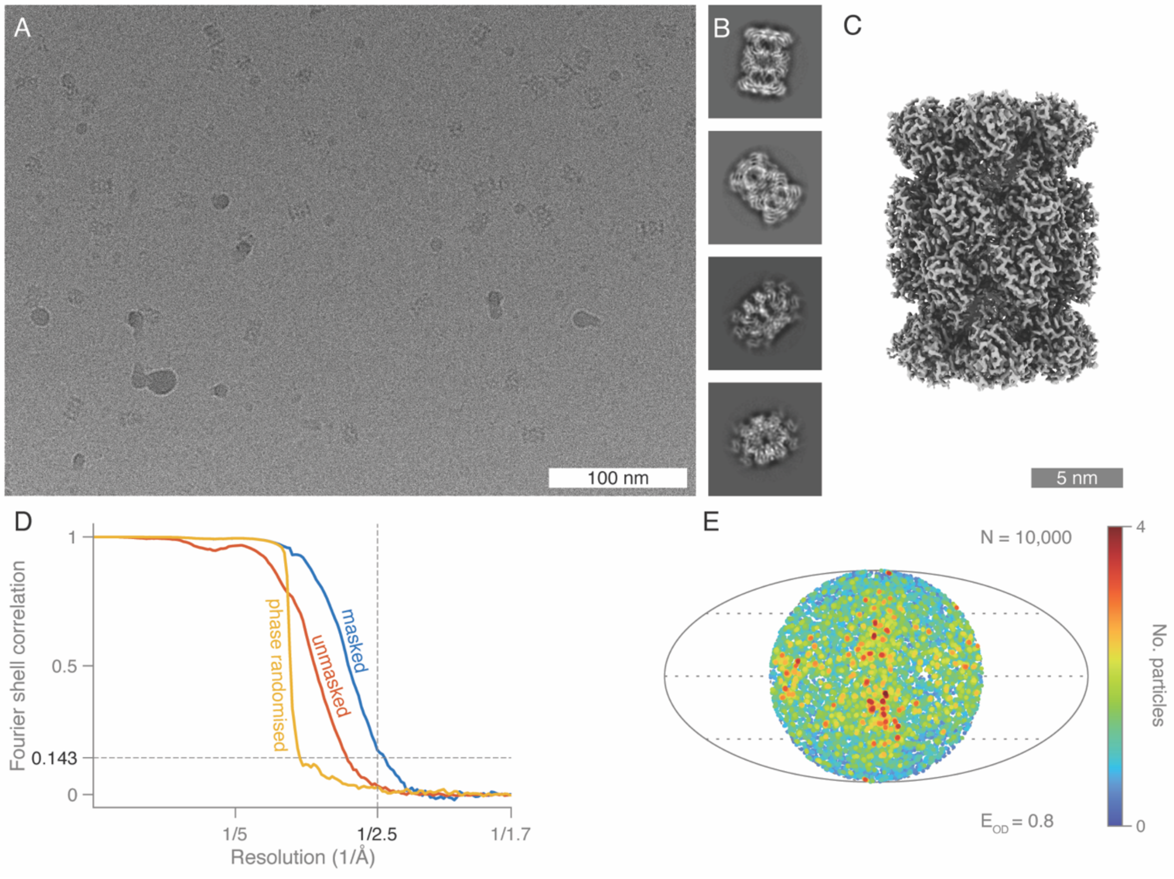


**Figure S8.** Structure determination of 20S proteasomes with 0.01% DDM. (A) Representative cryo-EM micrograph. (B) Four selected 2D class averages. (C) Final reconstructed cryo-EM 3D map. (D) Fourier shell correlation curves: unmasked dose-weighted half-maps (red), phase-randomised dose-weighted half-maps (yellow), and final, independently refined, masked, dose-weighted half-maps (blue). (E) Asymmetric particle orientation distribution plotted on a sphere, with the orientation distribution efficiency indicated.

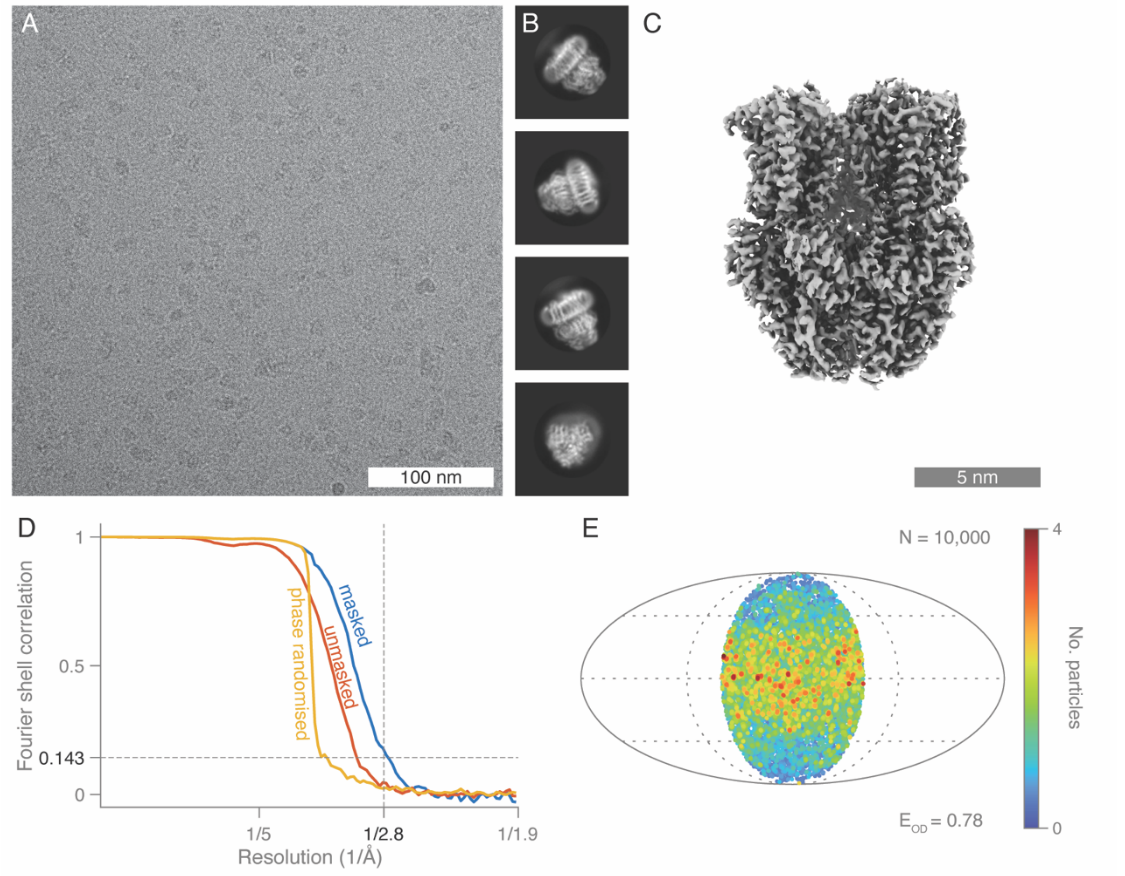


**Figure S9.** Structure determination of AcrB with 0.03% DDM. (A) Representative cryo-EM micrograph. (B) Four selected 2D class averages. (C) Final reconstructed cryo-EM 3D map. (D) Fourier shell correlation curves: unmasked dose-weighted half-maps (red), phase-randomised dose-weighted half-maps (yellow), and final independently refined, masked, dose-weighted half-maps (blue). (E) Asymmetric particle orientation distribution plotted on a sphere, with the orientation distribution efficiency indicated.

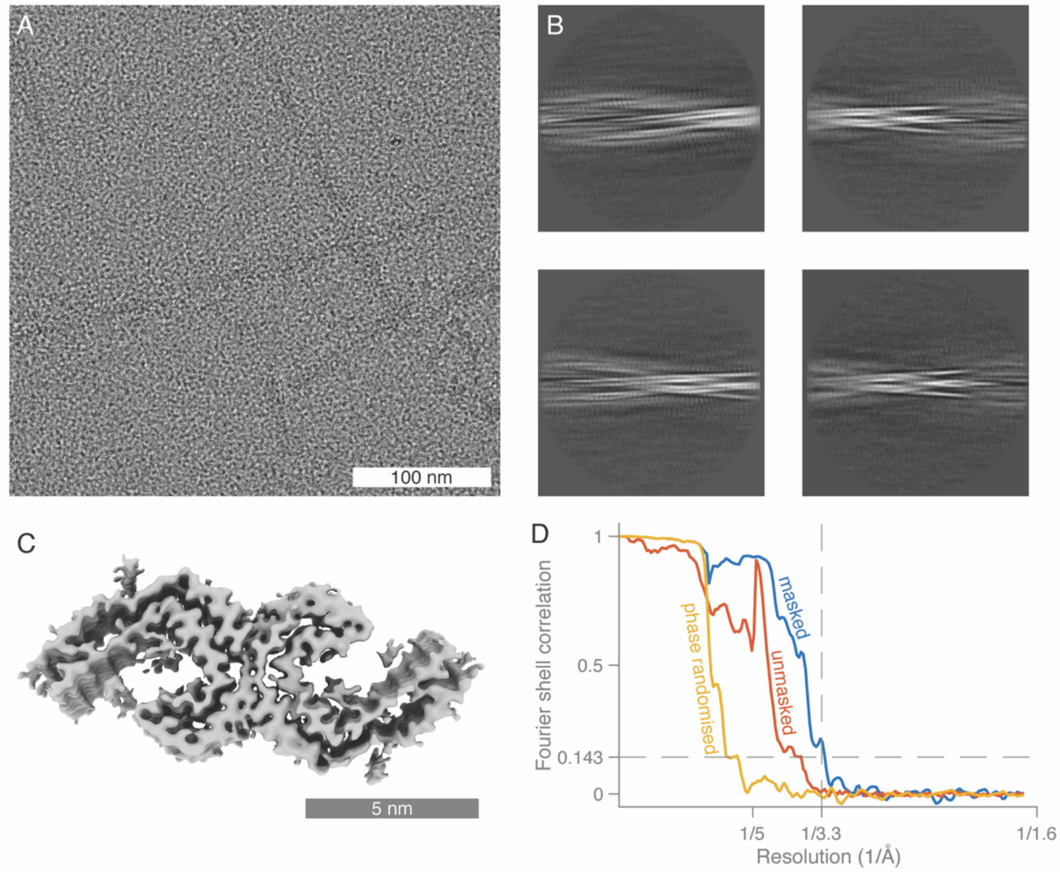


**Figure S10.** Structure determination of paired helical filaments. (A) Representative cryo-EM micrograph. (B) Four selected 2D class averages. (C) Final reconstructed cryo-EM 3D map. (D) Fourier shell correlation curves: unmasked dose-weighted half-maps (red), phase-randomised dose-weighted half-maps (yellow), and final independently refined, masked, dose-weighted half-maps (blue).

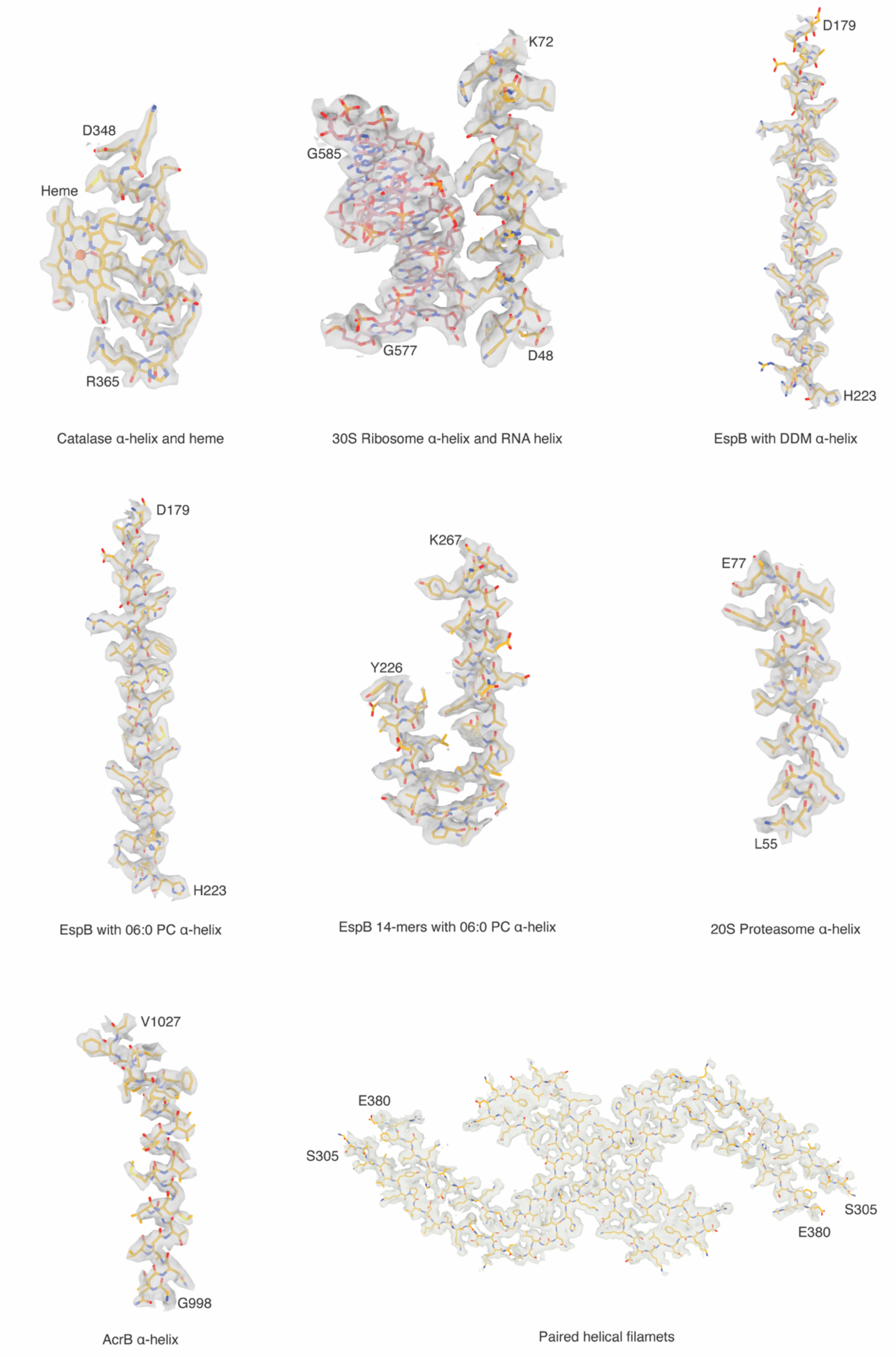


**Figure S11.** Cryo-EM map / atomic model superpositions for the structures prepared with foam film vitrification method.

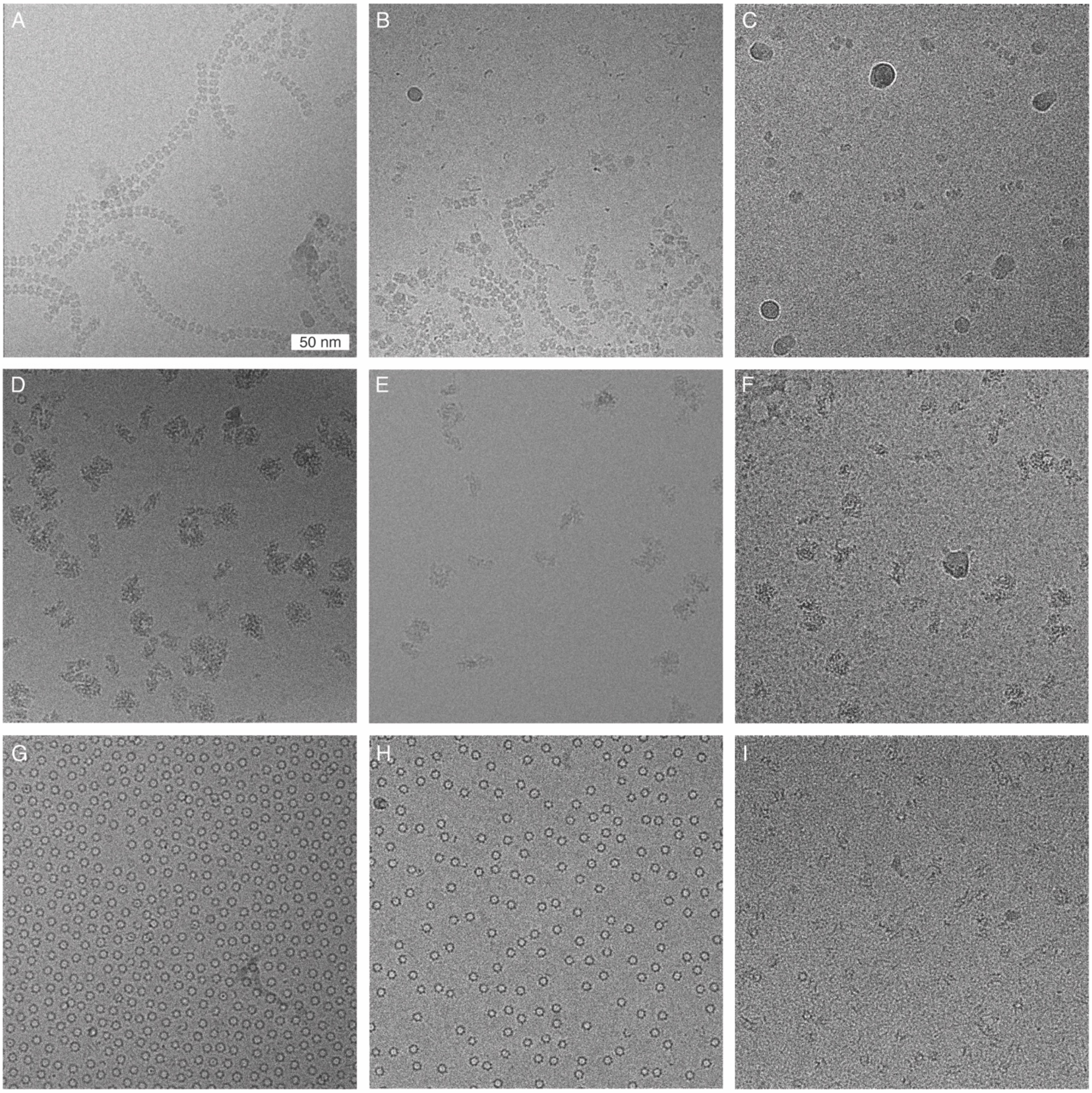


**Figure S12**. Example cryo-EM micrographs. (A) Catalase at a concentration of 0.1 g/L, prepared using the Vitrobot. (B) Catalase at a concentration of 0.5 g/L with 0.01% DDM, also prepared using the Vitrobot. (C) Catalase at a concentration of 3 g/L with 0.01% DDM, prepared using the foam film method. (D) 30S ribosomes at a concentration of 3.4 g/L, prepared using the Vitrobot. E) 30S ribosomes at a concentration of 3.4 g/L with 0.01% DDM, also prepared using the Vitrobot. (F) 30S ribosomes at a concentration of 3.4 g/L with 0.01% DDM, prepared using the foam film method. (G) EspB at a concentration of 1 g/L, prepared using the Vitrobot. (H) EspB at a concentration of 3 g/L with 0.01% DDM, also prepared using the Vitrobot. (I) EspB at a concentration of 5 g/L with 0.01% DDM, prepared using the foam film method.

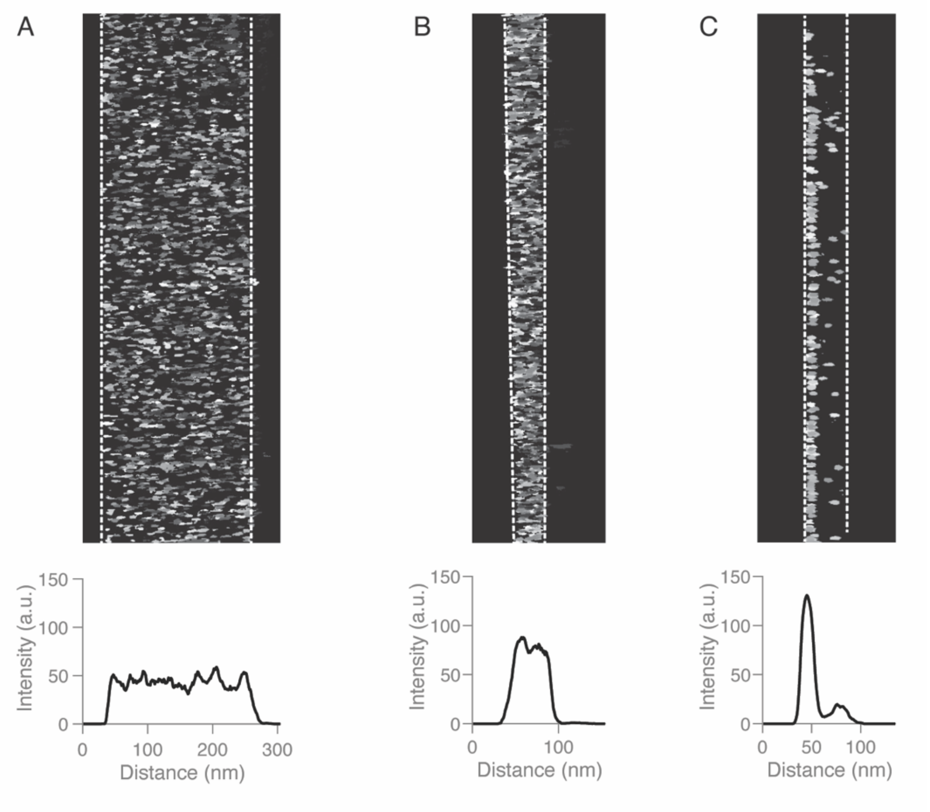


**Figure S13.** Distribution of iron-loaded ferritin particles in a cross section of vitrified ice as visualised by cryo-ET. The intensity plots display the average intensity along the vertical axis as a function of the horizontal direction. Iron cores from horse spleen ferritin are evenly distributed in both (A) thick and (B) thin samples prepared using the foam film method. (C) Iron particles in samples prepared using a Vitrobot accumulate more on the surface of the ice.

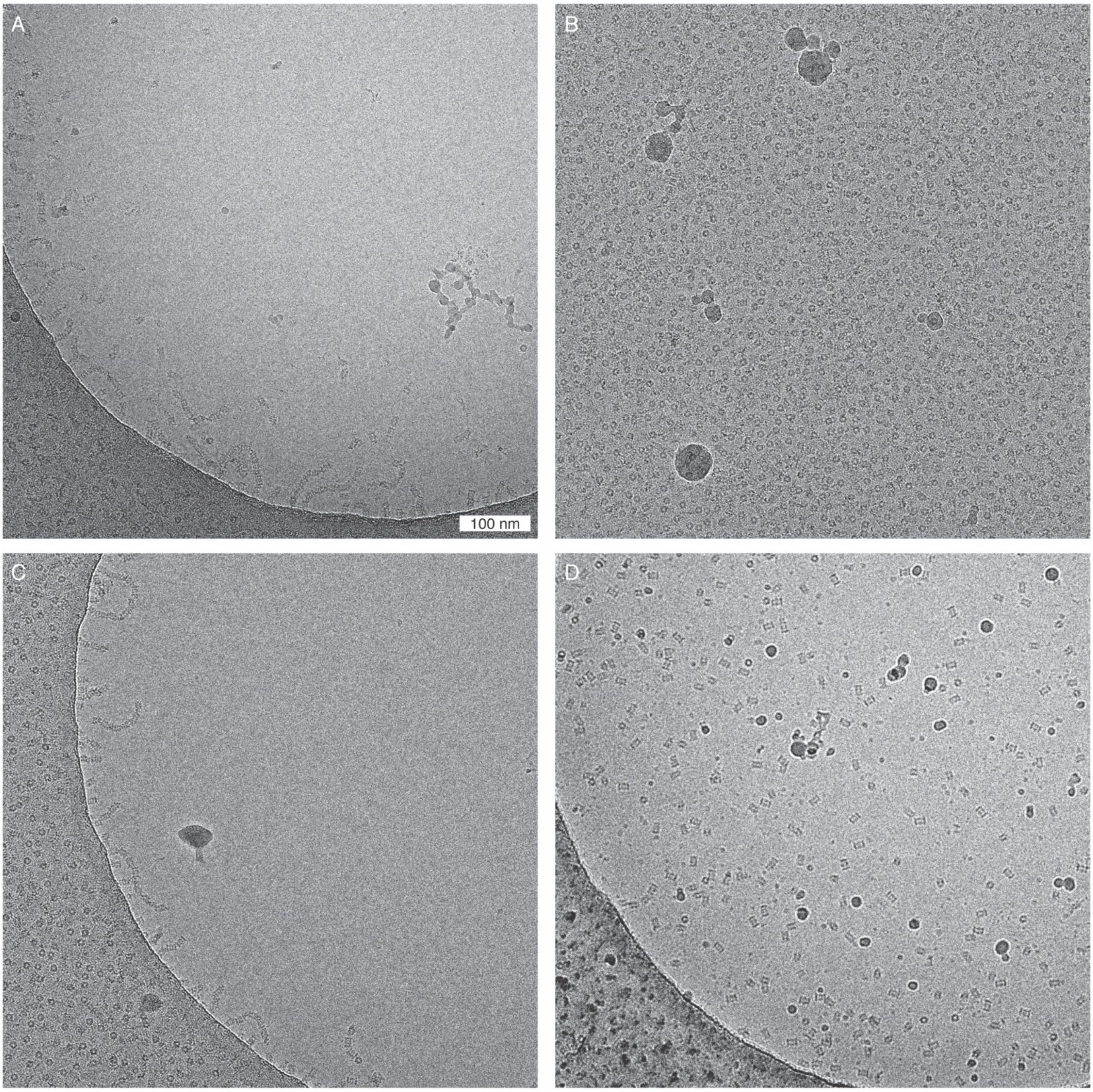


**Figure S14.** Example micrographs of 20S proteasomes, prepared using different methods. (A) 20S proteasomes at a concentration of 0.7 g/L, prepared with the Vitrobot, inside a foil hole. (B) 20S proteasomes at 0.7 g/L prepared with the Vitrobot on carbon foil. (C) 20S proteasomes at 0.7 g/L with 0.01% DDM and prepared with the Vitrobot. (D) 20S proteasomes at 0.7 g/L with 0.01% DDM, prepared using the foam film method.

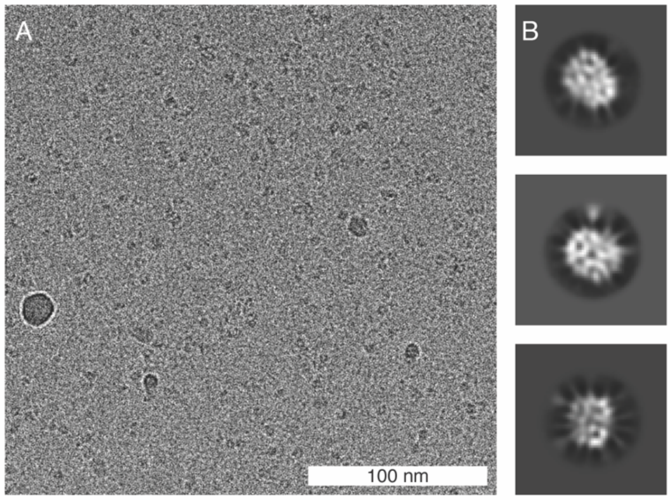


**Figure S15.** (A) Representative cryo-EM micrograph haemoglobin prepared using the foam film method. (B) Selected 2D class averages.

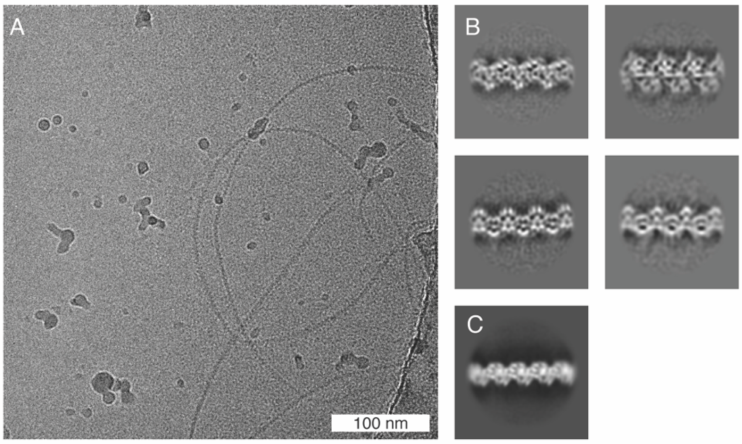


**Figure S16.** (A) Representative cryo-EM micrograph of FtsZ prepared using the foam film method. (B) Four selected 2D class averages showing seemingly different orientations. (C) The only 2D class from the dataset obtained from samples prepared with the Vitrobot.

**Table S1.** Cryo-EM data. Summary of cryo-EM data collections and data processing results for the samples prepared with the foam film cryo-EM vitrification method.

| | Catalase | 30S ribosomes | EspB | EspB 14-mers | EspB | 20S proteasomes | AcrB | Paired helical filament |
|---|---|---|---|---|---|---|---|---|
| Surfactant | DDM | DDM | DDM | 06:0 PC | | DDM | DDM | sarkosyl |
| Specimen support | Quantifoil Cu 300 mesh 1.2/1.3 | UltrAuFoil 300 mesh 1.2/1.3 | Quantifoil Cu 300 mesh 1.2/1.3 | Quantifoil Cu 300 mesh 1.2/1.3 | | Quantifoil Cu 300 mesh 1.2/1.3 | Quantifoil Cu 300 mesh 1.2/1.3 | UltrAuFoil 300 mesh 1.2/1.3 |
| Microscope | Krios | Krios | Krios | Krios | | Krios | Krios | Krios |
| Voltage (kV) | 300 | 300 | 300 | 300 | | 300 | 300 | 300 |
| Camera | Falcon 4i | Falcon 4i | Falcon 4i | Falcon 4i | | K3 | Falcon 4i | Falcon 4i |
| No. micrographs | 10,343 | 9,644 | 5,102 | 3,697 | | 5,212 | 8,828 | 3924 |
| No. picked particles | 1,105,755 | 778,822 | 1,091,474 | 1,047,418 | | 353,265 | 1,332,511 | 72976 |
| No. particles in reconstruction | 77,953 | 39,546 | 142,968 | 125,960 | 34,712 | 149,815 | 501,093 | 48999 |
| Nominal magnification | 96,000× | 96,000× | 165,000× | 130,000× | | 105,000× | 130,000× | 96,000× |
| Pixel size (A/pixel) | 0.824 | 0.824 | 0.744 | 0.955 | | 0.826 | 0.955 | 0.824 |
| Defocus range (um) | -0.6 to -2.4 | -0.6 to -2.4 | -0.6 to -2.4 | -0.6 to -2.4 | | -0.6 to -2.4 | -0.6 to -2.4 | -0.9 to -2.4 |
| Flux ($e^-/Å^2/s$) | 15.9 | 15.37 | 12.3 | 8.66 | | 21.5 | 9.11 | 13.8 |
| Fluence ($e^-/Å$) | 40 | 40 | 40 | 40 | | 40 | 40 | 40 |
| Fractions (no.) | 40 | 40 | 40 | 40 | | 40 | 40 | 40 |
| Symmetry imposed | D2 | C1 | C7 | D7 | C7 | C2 | C3 | Helical symmetry twist: -0.6, rise: 2.36 |
| Resolution (0.143 Fourier shell correlation, Å) | 2.6 | 3.45 | 2.72 | 2.34 | 3.07 | 2.45 | 2.77 | 3.28 |
| Efficiency of orientation distribution | 0.8 | 0.64 | 0.77 | 0.79 | 0.77 | 0.8 | 0.78 | - |
| B-factor ($Å^2$) | 140 | -96 | -152 | -105 | -88 | -100 | -118 | -100 |

#### Captions for Movies S1 to S15

**Movie S1**. Reconstructed tomograms of catalase at a concentration of 6 g/L with 0.01% DDM, prepared using the foam film method. Tilt series were recorded at nominal magnifications of ×53,000, with calibrated pixel sizes of 2.39 Å.

**Movie S2.** Reconstructed tomograms of catalase at a concentration of 0.1 g/L, prepared using the Vitrobot without surfactant. Tilt series were recorded at nominal magnifications of ×53,000, with calibrated pixel sizes of 2.39 Å.

**Movie S3.** Reconstructed tomograms of catalase at a concentration of 0.5 g/L with 0.01% DDM, prepared using the Vitrobot. Tilt series were recorded at nominal magnifications of ×53,000, with calibrated pixel sizes of 2.39 Å.

**Movie S4.** Reconstructed tomograms of 30S ribosomes at a concentration of 3.4 g/L with 0.01% DDM, prepared using foam film method. Tilt series were recorded at nominal magnifications of ×81,000, with calibrated pixel sizes of 1.514 Å.

**Movie S5.** Reconstructed tomograms of 30S ribosomes at a concentration of 3.4 g/L, prepared using the Vitrobot without surfactant. Tilt series were recorded at nominal magnifications of ×81,000, with calibrated pixel sizes of 1.514 Å.

**Movie S6**. Reconstructed tomograms of 30S ribosomes at a concentration of 3.4 g/L with 0.01% DDM, prepared using the Vitrobot. Tilt series were recorded at nominal magnifications of ×53,000, with calibrated pixel sizes of 2.39 Å.

**Movie S7**. Reconstructed tomograms of EspB at a concentration of 5 g/L with 0.01% DDM, prepared using foam film method. Tilt series were recorded at nominal magnifications of ×81,000, with calibrated pixel sizes of 1.514 Å.

**Movie S8.** Reconstructed tomograms of EspB at a concentration of 1 g/L, prepared using the Vitrobot without surfactant. Tilt series were recorded at nominal magnifications of ×81,000, with calibrated pixel sizes of 1.514 Å.

**Movie S9.** Reconstructed tomograms of EspB at a concentration of 3 g/L with 0.01% DDM, prepared using the Vitrobot. Tilt series were recorded at nominal magnifications of ×53,000, with calibrated pixel sizes of 2.39 Å.

**Movie S10.** Reconstructed tomograms of EspB 14-mers at a concentration of 2 g/L with 0.67% DDM, prepared using foam film method. Tilt series were recorded at nominal magnifications of ×81,000, with calibrated pixel sizes of 1.514 Å.

**Movie S11.** Reconstructed tomograms of EspB 14-mers at a concentration of 2 g/L with 0.67% DDM, prepared using the Vitrobot. Tilt series were recorded at nominal magnifications of ×53,000, with calibrated pixel sizes of 2.39 Å.

**Movie S12.** Reconstructed tomograms of 20S proteasomes at a concentration of 0.7 g/L with 0.01% DDM, prepared using foam film method. Tilt series were recorded at nominal magnifications of ×64,000, with calibrated pixel sizes of 1.38 Å.

**Movie S13.** Reconstructed tomograms of 20S proteasomes at a concentration of 0.7 g/L, prepared using the Vitrobot without surfactant. Tilt series were recorded at nominal magnifications of ×81,000, with calibrated pixel sizes of 1.514 Å.

**Movie S14.** Reconstructed tomograms of AcrB at a concentration of 3 g/L with 0.03% DDM, prepared using foam film method. Tilt series were recorded at nominal magnifications of ×81,000, with calibrated pixel sizes of 1.514 Å.

**Movie S15.** Reconstructed tomograms of AcrB at a concentration of 3 g/L with 0.03% DDM, prepared using the Vitrobot. Tilt series were recorded at nominal magnifications of ×53,000, with calibrated pixel sizes of 2.39 Å.
